# Locally balanced inhibition allows for robust learning of input-output associations in feedforward networks with Hebbian plasticity

**DOI:** 10.1101/2025.11.11.687798

**Authors:** Gloria Cecchini, Alex Roxin

## Abstract

In neural networks within the brain, the activity of a post-synaptic neuron is determined by the combined influence of many pre-synaptic neurons. This distributed processing enables mechanisms like Hebbian plasticity to associate sensory inputs with specific internal states, as seen in feedforward structures such as the CA1 region of the hippocampus. By modifying synaptic weights through Hebbian rules, sensory inputs can subsequently elicit outputs that consistently reflect their corresponding internal states. When input and output patterns are uncorrelated, this approach allows for the encoding of a large number of distinct associations, enabling efficient memory storage.

Our study demonstrates a critical limitation when output patterns become weakly correlated with input patterns through the intrinsic feedforward network’s connectivity. In these cases, the Hebbian rule preferentially strengthens synaptic weights shared across patterns, leading to a “freezing” of the network’s structure. This results in highly correlated output patterns over time, effectively reducing the network’s capacity to store diverse associations and limiting its flexibility in learning.

To address this challenge, we find that including a mechanism of locally balanced inhibition, which has been shown to be a key feature of cortical circuits in-vivo, counteracts the undesired correlations between inputs and outputs. By dynamically regulating inhibitory input, locally balanced inhibition prevents the over-strengthening of shared weights, restoring the network’s ability to maintain robust and flexible learning. This finding underscores the importance of inhibitory mechanisms in enabling efficient and adaptive information processing in neural circuits, offering insights into how biological networks maintain their remarkable capacity for associative learning.

## 1 Introduction

Associative learning in hippocampus and neocortex is often modeled with Hebbian plasticity, by which synapses strengthen between co-active pre- and post-synaptic neurons, and may weaken otherwise (Hebb (1949); Shatz (1992); Trappenberg (2002)). Learning through Hebbian mechanisms in recurrent networks has been the topic of intense study since the early 80s, e.g. Hopfield (1982); Amit et al. (1985); Tsodyks (1988); Amit and Fusi (1992); Amit and Brunel (1997); Battaglia and Treves (1998); Gilson et al. (2009); Zenke et al. (2015); Pereira and Brunel (2018); Feng and Brunel (2024). Such auto-associative networks store patterns as attractor states, which allow for pattern completion during recall given a partial cue. Learning in feedforward networks is generally modeled in one of two ways. First, unsupervised learning in feedforward networks with Hebbian plasticity can lead to the emergence of feature selectivity given sufficient competition between weights onto output units, either through normalization or recurrent connections Linsker (1986); Miller et al. (1989); Obermayer et al. (1992); Miller (1994). Alternatively, an association between imposed input and output patterns can be formed through a supervised rule such as stochastic gradient descent Rosenblatt (1958); Minsky and Papert (1969); Widrow and Stearns. In the latter case it remains unclear how such associations might actually be formed in the brain via local plasticity rules, e.g. Hebbian. Furthermore, the simplifying assumption of imposed input and output patterns, while reasonable from a normative standpoint, neglects the fact that outputs will be determined to some extent by the input pattern itself.

In this manuscript we study the process of memory storage in feedforward (hetero-associative) networks when the constraints of supervised learning and independent input- and output-patterns are relaxed. Specifically, we consider the case when such associations are formed solely through a local, Hebbian plasticity rule. Additionally, we consider the output pattern to be partially determined by the input pattern, namely we compute the postsynaptic output as the sum of the current synaptic drive from the target input and additional inputs, which can be approximated as a Gaussian noise term of adjustable amplitude. This design interpolates between two regimes: (i) strong input-output correlations when outputs depend only on the targeted input pathway, and (ii) vanishing input-output correlations when the influence of the targeted pathway on outputs is negligible. This interpolation allows us to isolate how input-output correlation shapes plasticity, network dynamics, and memory.

We show that when input-output coupling is strong (i.e., the postsynaptic output is dominated by the current synaptic drive), the network undergoes a *freezing transition*: successive readouts for uncorrelated inputs become highly similar. Mechanistically, binarized outputs preferentially activate high in-degree; Hebbian updates then potentiate those rows and depress low indegree rows, producing a progressive polarization of the connectivity (row-sum separation). Increasing the contribution of additional inputs weakens this correlation and recovers classical behavior, only when the influence of the targeted pathway on outputs is almost negligible. This raises the question: what mechanism preserves flexibility when output–input coupling cannot be neglected? Motivated by circuit evidence for inhibitory balance (Vogels et al. (2011); Renart et al. (2010); Zenke et al. (2015); Chistiakova et al. (2015); Bannon et al. (2020); Murphy and Miller (2009)), as well as in in-vivo recordings (Haider et al. (2006); Shu et al. (2003); Dehghani et al. (2016); Okun and Lampl (2008); Bhatia et al. (2019)), we introduce a locally balanced inhibition mechanism: each neuron receives an inhibitory term proportional to its own excitatory in-degree. When the inhibitory gain is tuned to the network’s activity level, the local balancing term effectively recenters postsynaptic drive, so output variability is determined by the current input rather than by pre-existing differences in synaptic in-degree. This breaks the bias toward repeatedly activating the same highly connected neurons, and consequently it decorrelates successive readouts. Moreover, in this regime we observe a clear improvement in memory strength.

All together, by introducing a mechanism of locally balanced inhibition, we demonstrate a simple yet powerful solution that restores flexibility and preserves memory capacity. These findings emphasize the importance of inhibitory regulation in maintaining the balance between stability and adaptability in neural learning systems.

## 2 Results

### 2.1 Dynamics of associative learning and interference in Hebbian networks

In this section, we introduce the main results and features of Hebbian plasticity as a mechanism for learning in feedforward neural networks (Hebb (1949); Shatz (1992); Trappenberg (2002)). First, Hebbian plasticity enables the formation of associations between input and output patterns. For instance, the presentation of a specific sensory cue, such as the sight of rain, can become associated with a behavioral response, like reaching for an umbrella (Fig. 1a). In a Hebbian framework, such associations are stored by modifying synaptic weights in a way that strengthens the connection between co-active pre- and post-synaptic neurons, and weakens it if one is inactive. Specifically, in this work, we consider the connectivity matrix *C*^*t*^ at time *t* to be binary. If 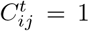, then the pre-synaptic neuron 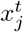 is connected to the post-synaptic neuron 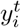. Moreover, we also consider the activity of the neurons to be binary, i.e., we focus only on the active/inactive status of each neuron, therefore 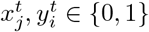. Hebbian plasticity follows a stochastic rule

**Fig. 1:**
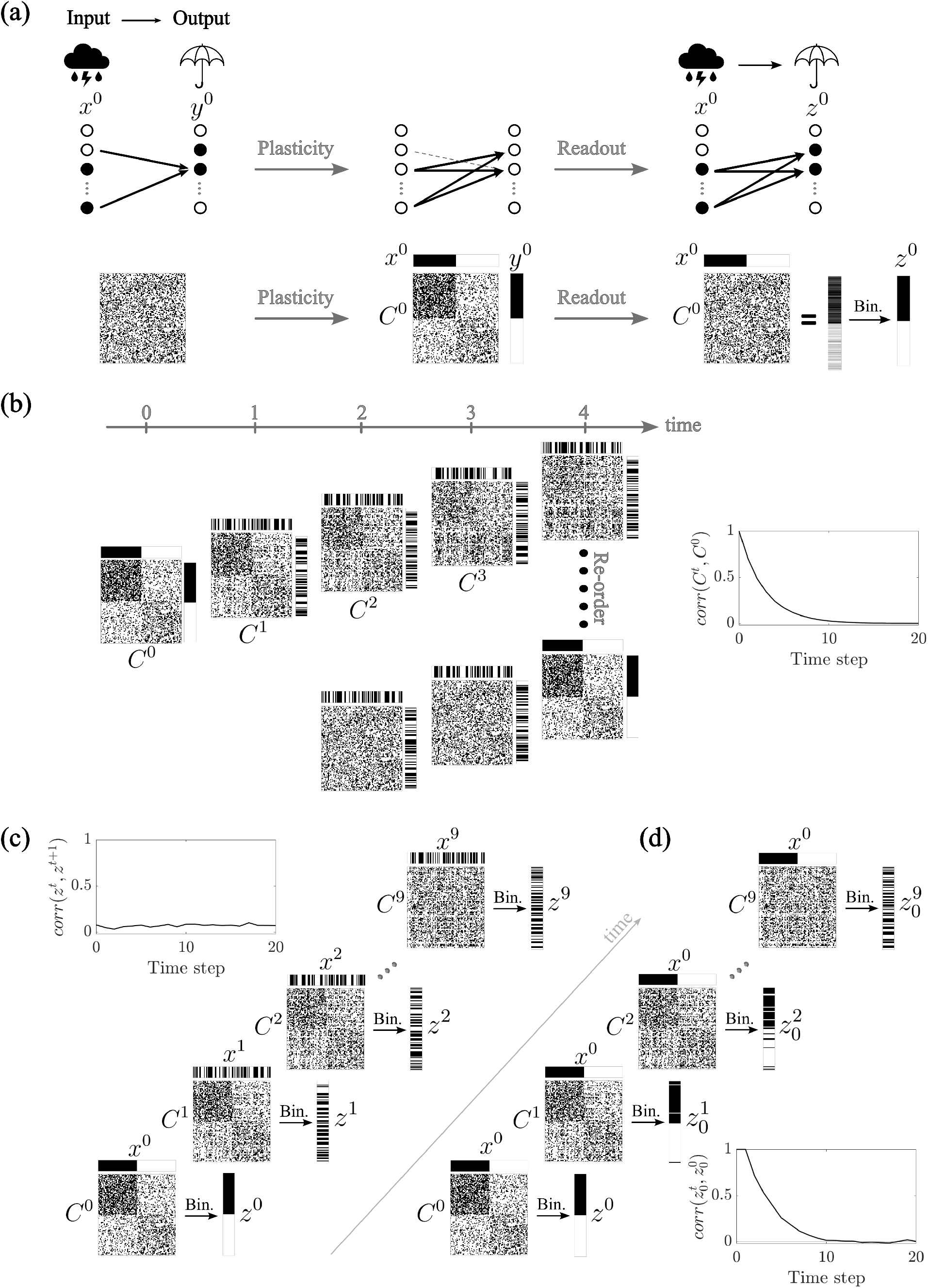
Dynamics of Associative Learning and Interference in Hebbian Networks. **(a)** Hebbian plasticity allows the association of input patterns (*x*^0^) with corresponding output patterns (*y*^0^) in a feedforward network. It does so by updating the connectivity matrix *C*, potentiating and depressing synapses depending on the state (active/inactive) of the respective connected cells. **(b)** As new (uncorrelated) associations are learned, the synaptic structure formed by previous associations is progressively overwritten, as shown by the decay in the correlation between the connectivity matrix *C* at time 0 and time *t*. The connectivity matrix is always correlated with the most recently stored patterns, as revealed by reordering the cells. **(c)** After learning, driven readouts (*z*^*t*^) remain largely uncorrelated, e.g. distinct sensory inputs lead to distinct outputs. The correlation between consecutive readouts *z*^*t*^ of random inputs *x*^*t*^ remains low. **(d)** Over time, the autocorrelation of an elicited readout 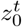 decays over time due to the overwriting from ongoing learning, leading to a progressive loss in recall fidelity. As a result, the correlation 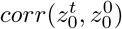 decreases and eventually approaches zero after a sufficient number of learning steps. – Parameters: *f* = 0.5, *p* = 0.4, *N* = 100 (*N* = 1000 for the correlation plots).

- if 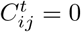 and 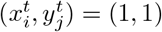, then 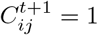 with probability *p*;
- if 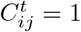 and 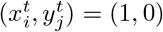 or (0, 1), then 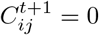 with probability *p*;
- if 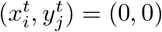, then 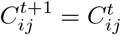.

Finally, the readout *z*^*t*^ is calculated as the multiplication of the connectivity matrix *C*^*t*^ and the input vector *x*^*t*^. In order to classify neurons as active or inactive, we set a threshold and binarize this result. The threshold value is chosen such that the fraction of active cell *f* remains constant across time. Once the network connectivity has reached a statistical steady-state, this threshold is simply a fixed value. All encoded patterns are taken to be random and uncorrelated unless otherwise noted.

As the network continues to learn new associations, previously formed synaptic structures are progressively altered (Fig. 1b). This is a property of networks with bounded synapses, and hence of biological networks Amit and Fusi (1992, 1994); Fusi and Abbott (2007); Fusi et al. (2021). Namely, when new learning drives a synapse to its bound, previously learned patterns become partially overwritten. As a consequence, memory traces decay smoothly in time and memory capacity is measured as the number of patterns encoded before this trace can no longer be detected. This is in contrast with the case of unbounded synapses, such as in the Hopfield network Hopfield (1982); Tsodyks (1988) where synapses are equally correlated with all stored patterns, and capacity is measured as the maximum number of patterns stored before all are lost simultaneously in the so-called blackout catastrophy Amit et al. (1985). Therefore, in networks with bounded synapses, unlike those with unbounded ones, ongoing learning is possible, but with the tradeoff of continual degradation of the memories over time.

This degradation due to overwriting is evident when examining the correlation *corr*(*C*^*t*^, *C*^0^) between the connectivity matrix *C* at the initial learning time (*t* = 0) and its state at a later time *t*. As new input-output pairs are encoded, the correlation between these matrices decays, reflecting the ongoing overwriting of old associations by new ones. Despite this turnover, the synaptic matrix remains consistently aligned with the most recently stored associations. This is demonstrated by reordering the neurons according to their current activity patterns: a clear structural organization emerges that reflects the latest learning episodes (Fig. 1b).

As shown in Fig. 1c, when the network is probed with novel learned (and uncorrelated) inputs, the resulting readouts *z*^*t*^ are distinct from one another. This ensures that different inputs elicit distinct outputs, an essential feature of reliable memory storage. However, the persistence of these associations is limited. Over time, as additional patterns are encoded, the output corresponding to an initially learned input pattern progressively changes (Fig. 1d). This is captured by the decay of the autocorrelation 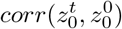, where 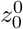 is the initial output elicited by an input *x*^0^, and 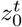 is the readout of the same input after *t* learning steps. The correlation decreases exponentially with each new association, eventually approaching zero, indicating a complete loss of recall fidelity. This phenomenon illustrates the inherent limitations of associative memory in ongoing learning in Hebbian networks, highlighting the trade-off between learning new information and retaining previously acquired associations.

### 2.2 An Input-Driven Framework for Hebbian Learning

As described in the previous section, hetero-associative learning assumes a given input- and output pattern. In order to study how such associations might arise in actual neuronal circuits, we must relax this constraint. Take, for example, the hippocampal CA1 region. Its activity is driven not only by input from CA3, but also by significant projections from other brain areas, such as the entorhinal cortex layer III (ECIII). If we wish to encode an association between a given input pattern from CA3 and a desired population response in CA1, we must take into account the relative influence of the CA3 inputs on the output pattern, compared to other pathways.

To do this, we compute the output as the result of the total input: that of the specific input pattern (e.g., from CA3) which we wish to associate with the output pattern, and additional inputs representing the influence of other brain areas (e.g., ECIII), neuromodulatory inputs or any change in excitability not linked to the input pathway of interest, see Fig. 2a. Mathematically, we model the output *y*^*t*^ as the result of the sum of the target input *C*^*t*−1^*x*^*t*^ and additional inputs 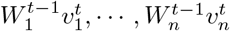, see Fig. 2a. Assuming input from the non-targeted pathways are uncorrelated with the one of interest, the sum of these additional inputs is equivalent to Gaussian noise *σξ* with *ξ* ~ *N* (0, 1). The parametrization of the standard deviation *σ* allows us to modulate its relative influence on the output. Therefore, the output is calculated as 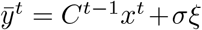 (Fig. 2b). In order to classify neurons as active or inactive, we set a threshold and binarize 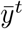. The threshold value is chosen to ensure a particular coding fraction *f*. Once the binarized output *y*^*t*^ is calculated, the network undergoes plasticity and the connectivity matrix is updated as described in the previous section, resulting in *C*^*t*^. Finally, the readout *z*^*t*^ is driven by the target input *x*^*t*^ and the corresponding connectivity matrix *C*^*t*^ (Fig. 2c-d). Specifically, the readout *z*^*t*^ is the binarization of 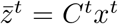, again keeping constant the fraction of active cell *f*.

**Fig. 2:**
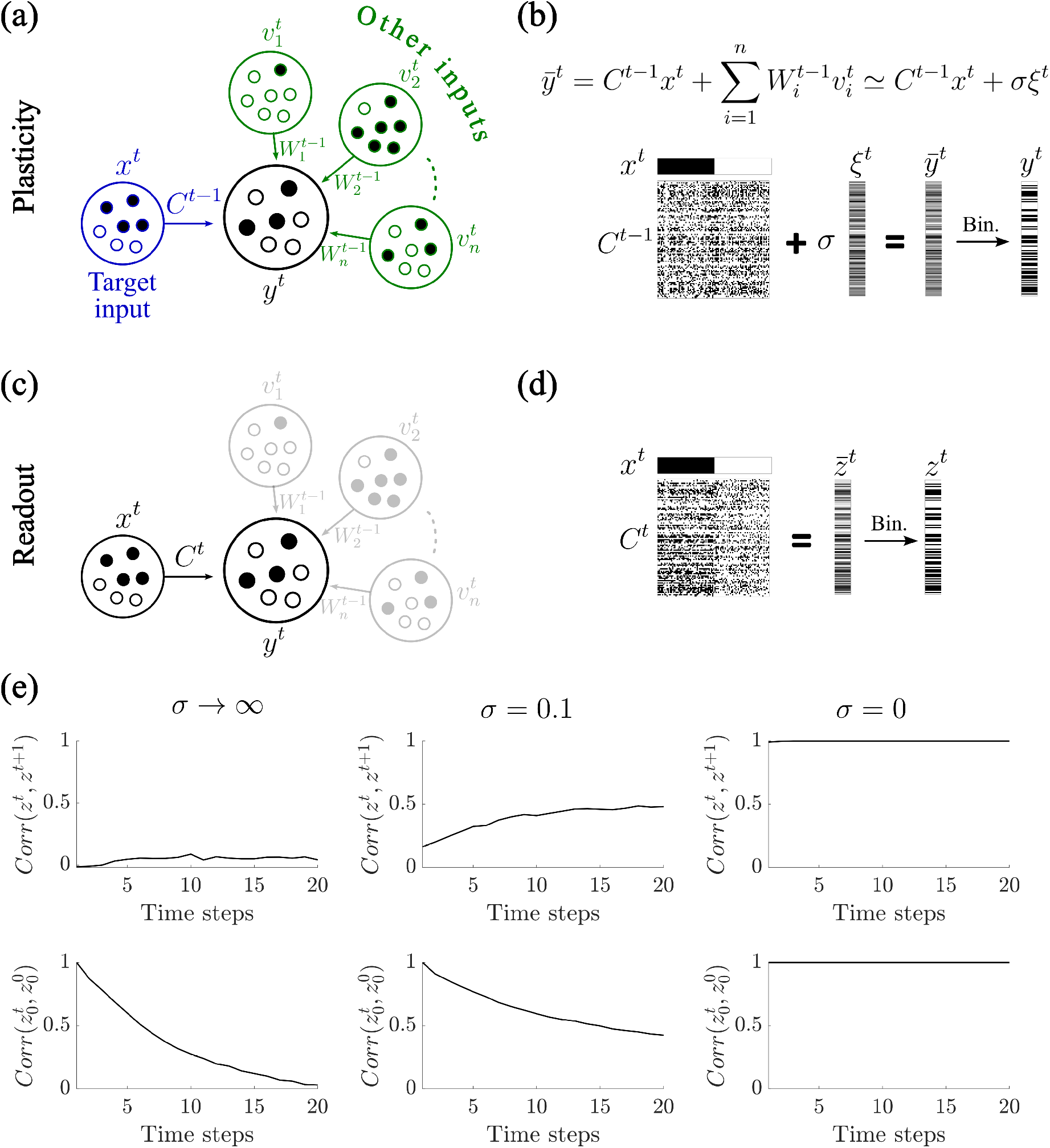
An Input-Driven Framework for Hebbian Learning. **(a-b)** The output *y*^*t*^ reflects the sum of a target input pattern *x*^*t*^ and additional inputs 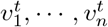, resulting in partial input-output correlations. These additional inputs 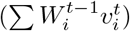 can be modeled as Gaussian noise (*σξ*). The output *y*^t^ is the binarization of 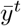 that is obtained as the sum of the contributions from the target input *C*^*t*−1^*x*^*t*^ and the additional inputs. **(c-d)** The readout *z*^*t*^ is driven by the target input pattern *x*^*t*^ and the corresponding connectivity matrix *C*^*t*^, after plasticity. **(e)** Correlation of neighboring readouts *corr*(*z*^*t*^, *z*^*t*+1^) and auto-correlation 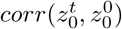 elicited by an input *x*^0^. For *σ* → ∞, we have the same scenario as in Fig. 1. For *σ* = 0.1, i.e., in the presence of weak correlation between output and synaptic input strength, the network produces readouts *z*^*t*^ that remain strongly similar across time. For *σ* = 0, the readouts *z*^*t*^ become identical. – Parameters: (b,d) *N* = 100, *f* = 0.5, *p* = 0.4, *σ* = 0.1 (e) *N* = 1000, *f* = 0.5, *p* = 0.2.

To investigate how the partial correlation between output and synaptic input strength influences network plasticity, we varied the strength of the non-target input pathways relative to the target input pathway, i.e., *σ*. This manipulation simulates a spectrum of scenarios, ranging from fully input-driven outputs (low *σ*) to outputs dominated by the other inputs (high *σ*), see Fig. 2e. Our simulations reveal that even when the correlation between output and synaptic input is weak the network exhibits a pathological dynamic: successive readouts *z*^*t*^ remain highly similar over time. In the extreme case where the output is entirely determined by the structured input (*σ* = 0), all readouts *z*^*t*^ become identical. This leads to a complete loss of learning flexibility, as the network effectively freezes into a fixed response. In contrast, for large *σ* values, the output is no longer systematically biased by the synaptic input structure, thereby recovering the conditions assumed in classical Hebbian paradigms. In this regime, the network behaves as previously described in Fig. 1, where uncorrelated inputs give rise to distinct outputs, enabling robust associative learning.

These results underscore the critical role of output-input correlations in shaping the long-term plasticity of feedforward networks and highlight the potential stabilizing role of additional inputs in maintaining learning flexibility. In the next section, we perform a more systematic analysis on the impact of an output that is correlated to the synaptic input strength, on network plasticity.

### 2.3 Correlation between output and synaptic input strength impairs associative learning

In the previous section, we presented some illustrative examples demonstrating how correlation between output activity and synaptic input strength can profoundly affect the dynamics of Hebbian plasticity in feedforward networks. In this section, we extend our analysis with a more systematic and quantitative investigation, aimed at characterizing the network’s plastic behavior across a continuum of input-output correlation strengths. To this end, we varied the amplitude *σ* of the noise term, which modulates the extent to which the output is driven by the target input versus other inputs. Small values of *σ* correspond to outputs tightly determined by the synaptic input, whereas large values of *σ* simulate conditions in which outputs are dominated by additional, non-specific inputs.

To evaluate the effects of *σ* on the network’s learning dynamics, we focus on the correlation 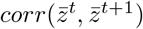 between consecutive readouts before binarization 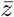, as a metric to quantify the degree of freezing in the network’s output. Since we consider uncorrelated inputs, high correlations between successive readouts indicate reduced flexibility and a tendency toward rigid, self-reinforcing representations.

We first analyze the case of *σ* = 0, where the output is entirely determined by the target input. As shown in Fig. 3a, in this regime the correlation between consecutive readouts progressively increases and eventually converges to 1, regardless of other network parameters such as the sparsity of activation *f* or the synaptic plasticity probability *p*. This convergence means that, over time, the network settles into a fixed readout pattern that is insensitive to the input identity, indicating a complete loss of representational flexibility. Interestingly, this convergence follows a power-law trajectory, as illustrated in Fig. 3b. We extracted the power-law exponent *b* across different values of *f* and *p* and found that both parameters modulate the rate at which this rigidity emerges: higher values of *f* or *p* accelerate the convergence (Fig. 3c). The asymptote value *c* was 1 for any combination of *f* and *p*.

**Fig. 3:**
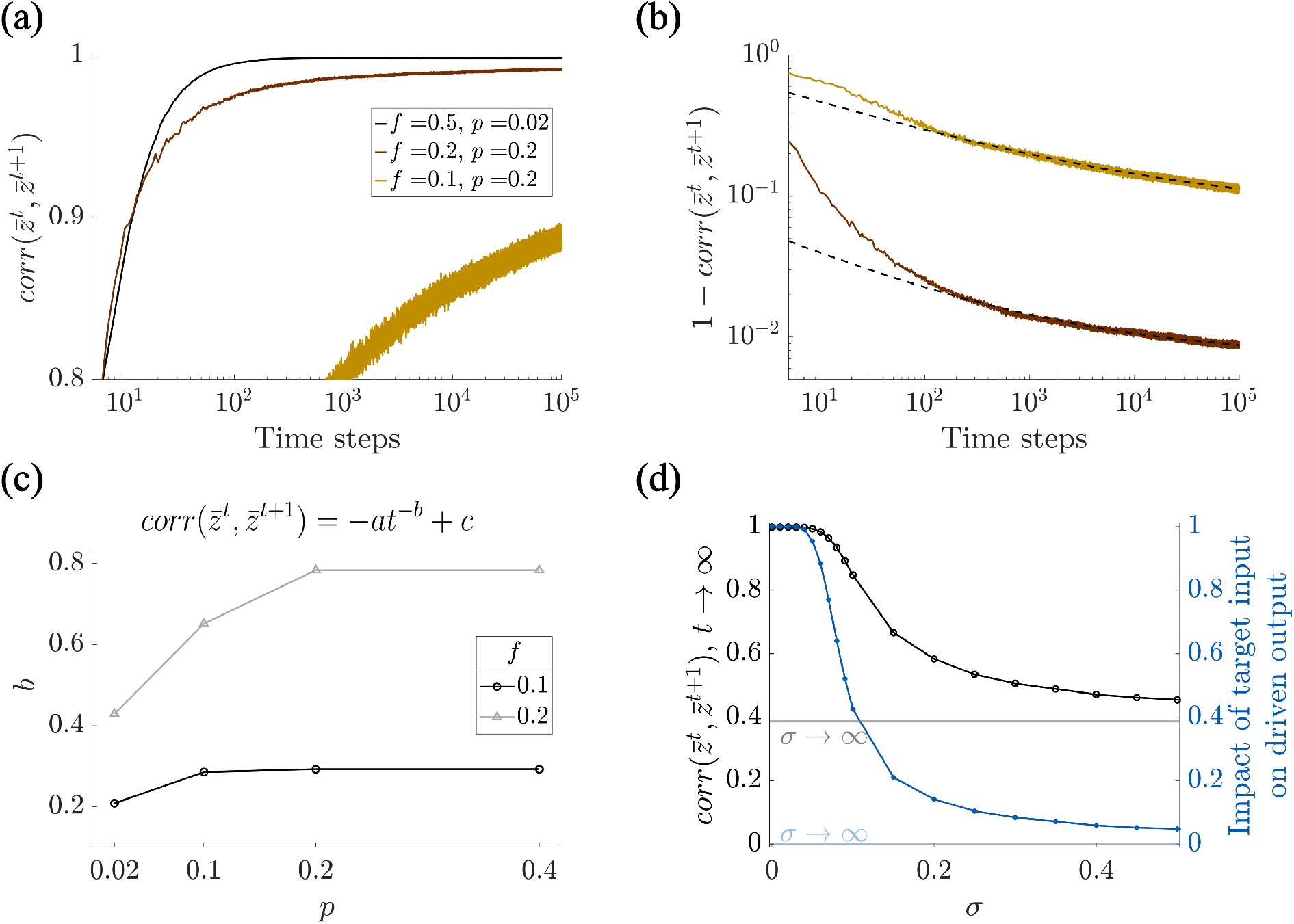
Correlation between output and synaptic input strength impairs associative learning. **(a)** Uncorrelated inputs yield identical readouts 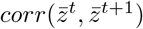 goes to 1) after learning when the effect of other input pathways is zero (i.e. *σ* = 0), for any fraction of active cells *f* and probability of potentiation/depression *p*. **(b)** The correlation 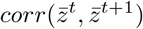 converges to one following a power-law (dashed lines: power-law fit). **(c)** The extracted power-law parameters for the power (*b*) reveal the dynamics of readout convergence (*c* = 1 for any *f* and *p*). **(d)** (Black) The influence of other inputs prevents convergence of the correlation 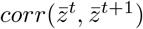, which decreases with increasing *σ*. (Blue) Impact of target input on the driven output, calculated as *corr*(*C*^*t*−1^*x*^*t*^ + *σξ*^*t*^, *C*^*t*−1^*x*^*t*^). – Each curve shows the mean over 50 simulations. Parame-ters: *N* = 1000, (a-c): *σ* = 0, (d): *f* = 0.5, *p* = 0.2.

Next, we explored how increasing the amplitude of the noise term *σ*, representing an increasing contribution of other inputs, affects the network’s tendency toward freezing. We systematically varied *σ* and measured the asymptotic value of 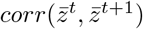, both numerically and via power-law fitting. The results, summarized in Fig. 3d (black), show a clear trend: as *σ* increases, the correlation between successive readouts decreases monotonically. These findings imply that additional inputs can prevent the network from entering a fully frozen state, thereby preserving its ability to encode diverse patterns.

Finally, we quantified the impact of the additional inputs on the driven output, specifically, how much the purely driven output *C*^*t*−1^*x*^*t*^ differs from the total output *C*^*t*−1^*x*^*t*^ + *σξ*^*t*^ as a function of the noise amplitude *σ*. To this end, we computed the correlation between these two signals. i.e., *corr*(*C*^*t*−1^*x*^*t*^ + *σξ*^*t*^, *C*^*t*−1^*x*^*t*^), which serves as a measure of the impact of the target input on the driven output. As shown in Fig. 3d (blue), increasing the noise leads to a progressive decorrelation between the two, as expected. However, a comparison with the correlation 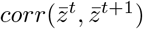 (in black) reveals a more nuanced picture: to fully decorrelate the output from the intrinsic bias of the network, the noise must be strong enough to almost completely override the contribution of the target input. In other words, mitigating network freezing through noise alone requires driving the output toward a nearly random pattern, essentially reverting to the classical Hebbian learning scenario in which outputs are externally imposed. Since the aim of this study is to develop a model in which outputs are determined by a target input, we must seek an alternative solution to this problem.

Although not shown here, we verified that once the system has reached a stationary state, i.e. enough patterns have been encoded so that the network statistics are stable, there is no need for a dynamic threshold to ensure a stable coding fraction *f* for a given value of *σ*. That is, if the network works at capacity, then everything else being the same, the coding fraction is expected to be constant in time, when setting a fixed threshold. In other words, simulations with a fixed threshold do not alter any of the conclusions of the paper.

In the next section, we investigate in greater detail the mechanisms underlying network freezing, particularly in scenarios where the output remains strongly correlated with the synaptic input strength.

### 2.4 Mechanism of network freezing due to high correlation between output and synaptic input strength

To understand the emergence of rigid dynamics in a feedforward network under Hebbian plasticity, we examined the consequences of strong correlations between output activity and synaptic input strength. Intuitively, once a neuron loses enough synapses, its input drops below threshold and it falls silent. Because the plasticity rule potentiates only co-active pairs, a silent postsynaptic neuron cannot regain synapses and instead tends to undergo further depression, effectively freezing it in a no-input state. Conversely, a neuron with sufficiently many synapses keeps firing, and co-activity sustains its potentiated inputs.

In Fig. 4a, we begin with a randomly initialized synaptic matrix. For visualization purposes, we reorder the rows of the matrix such that the resulting output vector 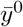 exhibits a descending gradient of values. This reordering allows us to visualize, in later steps, how Hebbian learning affects different subsets of the network. The distribution of output entries in 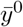 is shown using a histogram, where the first part of the vector is depicted in black and the second part in gray. In order to classify neurons as active or inactive, we set a threshold and binarize 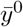. By construction, the binarization of this graded output yields a binary pattern *y*^0^, with the first part consisting entirely of ones and the second part of zeros. This binarized output pattern *y*^0^ is then used to do Hebbian plasticity together with the input *x*^0^, as explained in Sec. 2.1.

**Fig. 4:**
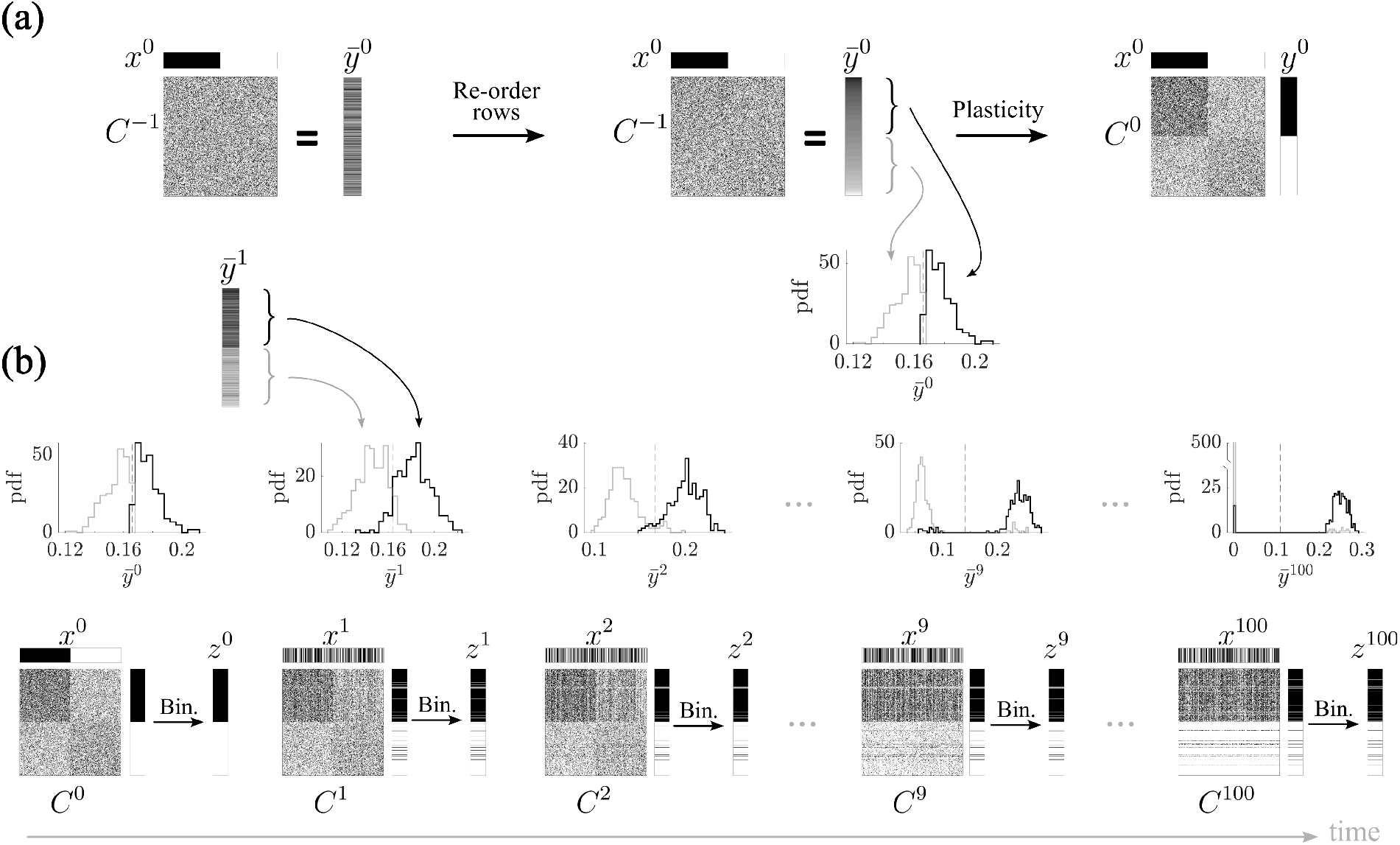
Mechanism of network freezing due to high correlation between output and synaptic input strength. **(a)** Starting from a random synaptic matrix (*C*^−1^), the rows are reordered (only at this initial step) to produce an output vector 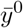 with values arranged in decreasing order. The histogram of the first part of its entries is shown in black, and the second one in gray. Binarizing 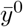 yields the vector *y*^0^, where the first part consists of 1s and the second one of 0s. Following Hebbian plasticity, the corresponding synaptic matrix is updated (*C*^0^), by potentiation and depression with probability *p*. **(b)** In the subsequent steps, the two parts of the output 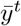 (without further re-ordering the rows) become increasingly separated. As a result, the binarized vector *y*^*t*^ consists mainly of 1s in the first part and 0s in the second one, further reinforcing the synaptic structure established in the previous step and amplifying the network’s rigidity. As the output entries become increasingly separated, the synaptic matrix *C*^*t*^ develops a more pronounced structure, with potentiated and depressed rows clearly diverging. This growing separation causes the network to produce identical readouts *z*^*t*^ over time. – Parameters: *N* = 500, *f* = 0.5, *p* = 0.2, *σ* = 0.

In subsequent learning steps, the asymmetry of the 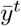 entries becomes amplified (Fig. 4b). As the network continues to process new inputs and undergo plasticity, already at time *t* = 1, the output vector 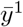 exhibits an even stronger separation between the first and second parts, as shown by the histograms. Consequently, its binarized version *y*^1^ remains almost identical to *y*^0^. This pattern of activity leads to further strengthening and weakening of the same subsets of synapses, reinforcing the structure already imprinted in the matrix during previous steps. Over time, the network’s synaptic matrix becomes increasingly polarized, with a clear division between potentiated and depressed rows (see *C*^100^ in Fig. 4b).

This progressive separation of output activity and its corresponding synaptic changes establishes a positive feedback loop: once a pattern of activity becomes dominant, it increasingly biases future outputs and plasticity events toward reinforcing the same structure. As a result, the network enters a frozen state in which the output becomes insensitive to new inputs. After a few time steps, regardless of the input presented, the network produces identical readouts. This collapse of output diversity reflects a pathological rigidity in the network’s dynamics, fundamentally impairing its capacity for associative learning and memory storage.

In the next section, we build upon the insights gained from the analysis of network freezing dynamics. Specifically, we aim to develop and investigate a biologically inspired mechanism capable of counteracting the onset of rigidity in network dynamics. By introducing a targeted intervention, our goal is to preserve the network’s capacity for flexible associative learning while mitigating the detrimental effects of excessive input-driven reinforcement.

### 2.5 Locally balanced inhibition counteracts input-driven rigidity

As shown in the previous sections, in feedforward networks governed by Hebbian plasticity, output activity that is exclusively determined by excitatory input can lead to highly stereotyped and inflexible activation patterns. In such scenarios, neurons with a large excitatory synaptic input are consistently activated, while those with fewer inputs are persistently suppressed. This biased pattern of activity reinforces the same subset of neurons across learning episodes, ultimately leading to network rigidity and reduced capacity for storing diverse associations.

To address this issue, we introduce a mechanism of locally balanced inhibition, which has been shown to be a key feature of cortical circuits both theoretically (Vogels et al. (2011); Renart et al. (2010); Vreeswijk and Sompolinsky (1996); Murphy and Miller (2009); Rubin et al. (2015); Koch et al. (2010)), and in-vivo (Haider et al. (2006); Shu et al. (2003); Dehghani et al. (2016); Okun and Lampl (2008); Murphy and Miller (2009); Ozeki et al. (2009)). In our model, we model locally balanced inhibition by making the inhibitory input to each neuron proportional to its excitatory synaptic input (Fig. 5a-b). Specifically, the output at time *t* is the binarization of

**Fig. 5:**
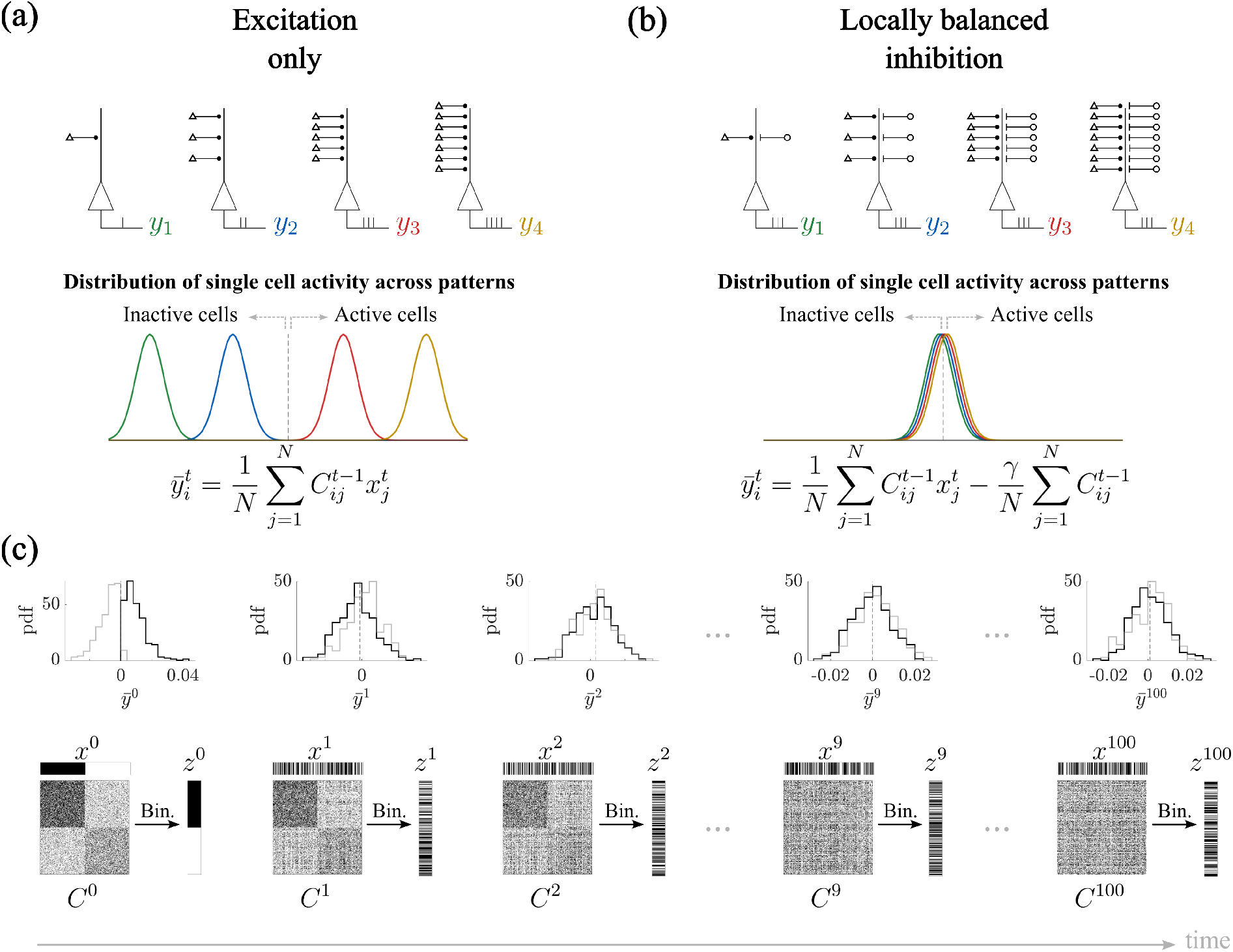
Locally balanced inhibition counteracts input-driven rigidity. **(a)** When output activity is solely driven by excitatory input, neurons (*y*_*i*_) with high in-degree are consistently selected as active, while those with low in-degree remain inactive, leading to biased activation patterns. The colored curves show a schematic of the distributions of the entries of *y*_*i*_, calculated as 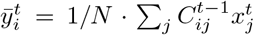. **(b)** Each output neuron *y*_*i*_ receives inhibitory input scaled to its total excitatory input in-degree 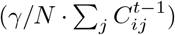, providing a neuron-specific balancing mechanism. **(c)** This local inhibitory balancing prevents progressive separation of output values 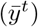, thereby avoiding matrix freezing and preserving the network’s capacity for flexible learning (compare to Fig.4). – Parameters: *N* = 500, *f* = 0.5, *p* = 0.4, *γ* = 0.5.

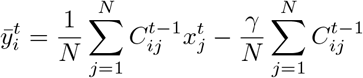

where *N* is the number of neurons, and *γ* is the inhibitory constant. This neuron-specific inhibitory feedback effectively normalizes the total synaptic drive each cell experiences, thereby eliminating the competitive advantage conferred by high in-degree. By counteracting the dominance of highly connected neurons, this inhibitory balancing mechanism prevents the progressive separation of output values that would otherwise emerge during repeated learning (compare Fig. 5c with Fig.4b).

The inhibitory mechanism just described enables robust learning of input-output associations only when appropriately tuned. Fig. 6a shows how the correlation *corr*(*z*^*t*^, *z*^*t*+*τ*^) between neighboring readouts evolves by changing the inhibitory constant *γ*; for this figure we set the fraction of active cells to *f* = 0.5. For *γ* not large enough (*γ <* 0.5), the inhibitory feedback is not able to counterbalance the synaptic drive, and therefore, eventually, the system converges into a rigid state, i.e., *corr*(*z*^*t*^, *z*^*t*+1^) = 1. On the other hand, if the inhibition is too strong (*γ >* 0.5), the correlation becomes negative, i.e., cells with low in-degree are first boosted and then strongly depressed at the following time step, causing *corr*(*z*^*t*^, *z*^*t*+1^) → −1 and *corr*(*z*^*t*^, *z*^*t*+2^) →1 for *γ >>* 0.5. When appropriately tuned (*γ* = 0.5), the network retains the ability to generate distinct and reliable output patterns in response to different input stimuli, i.e., *corr*(*z*^*t*^, *z*^*t*+*τ*^) = 0 for every *τ*.

**Fig. 6:**
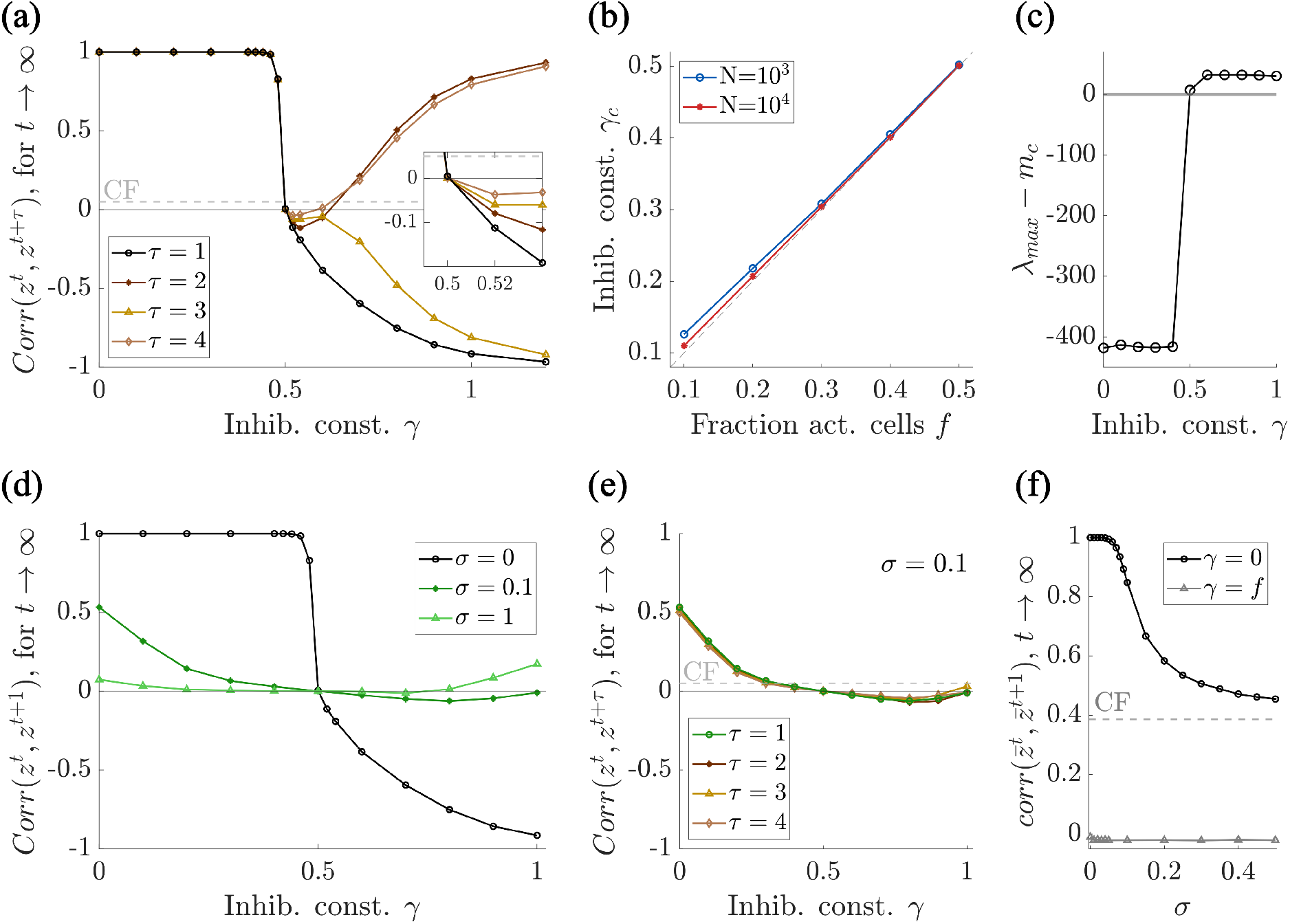
Locally balanced inhibition enables robust learning of input-output associations. **(a)** Appropriate levels of inhibition (*γ* = 0.5) restore the network’s ability to form reliable input-output associations, as shown by the correlation of the first (*τ* = 1) and subsequent (*τ* = 2, 3, 4) nearest neighbor (i.e. *corr*(*z*^*t*^, *z*^*t*+*τ*^) = 0 for *γ* = 0.5). **(b)** The inhibitory constant *γ* that minimize, in absolute value, the correlation *corr*(*z*^*t*^, *z*^*t*+1^) between neighboring readouts (called *γ*_*c*_) is proportional to the fraction of active cells *f*. **(c)** The rescaled largest eigenvalue vanishes only when *γ* = *f*. **(d)** As in panel (a), for *γ* = *f* = 0.5, the correlation *corr*(*z*^*t*^, *z*^*t*+1^) becomes zero, also when additional inputs are considered, i.e., for *σ* = 0.1, 1. **(e)** For *σ* = 0.1, the correlation of any subsequent neighbor *corr*(*z*^*t*^, *z*^*t*+*τ*^) becomes zero for *γ* = *f*. **(f)** When setting *γ* = *f*, the correlation *corr*(*z*^*t*^, *z*^*t*+1^) is zero for any value of *σ* (black line is the same as in Fig. 3d). – Each curve shows the mean over 10 simulations. Parameters: *p* = 0.2, *f* = 0.5, *N* = 5000.

We investigated if changing the parameters for the fraction of active cells *f* alters the trend of the correlation *corr*(*z*^*t*^, *z*^*t*+1^) as a function of *γ*. We found that the curve profile is qualitatively the same, but the value of *γ* that returns *corr*(*z*^*t*^, *z*^*t*+1^) = 0, called *γ*_*c*_, equals *f*, see Fig. 6b.

In order to have an intuition of why setting *γ* = *f* leads to *corr*(*z*^*t*^, *z*^*t*+1^) = 0, we calculate the mean of the output 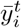

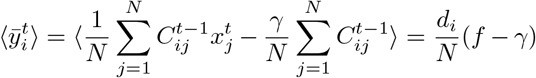

where *d*_*i*_ is the in-degree of each neuron *i*. We noticed that for *γ* = *f*, then 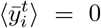, which means that the variability due to the heterogeneity in mean in-degrees is eliminated, and therefore strong/weak synapses will not necessarily be strengthened/weakened even more.

As an additional demonstration that *γ* = *f* restores full flexibility in the network, we calculated the eigenvalues of the connectivity matrix (*C*) after the system reached a steady state (*t* = 10^3^). For every value of *γ*, the eigenvalues are distributed within a disk centered at zero, except for one outlier lying on the real axis at varying distances depending on *γ* (see studies on non-Hermitian (Ginibre) matrices, e.g., O’Rourke and Renfrew (2013); Girko (1985)). In particular, this outlier, i.e., the largest eigenvalue *λ*_*max*_, scales as *N* ⟨*C*⟩. To assess the role of this outlier, we calculated the theoretical mean *m*_*c*_ expected if the network had evolved according to the classical framework (Sec. 2.1). This value was derived in Devalle et al. (2025), see supplementary materials. We then plotted the distance of the outlier eigen-value *λ*_*max*_ from the expected scaled mean *Nm*_*c*_ as a function of *γ*, shown in Fig. 6c for *f* = 0.5.

We observed that *λ*_max_ −*m*_*c*_ = 0 only when *γ* = *f* = 0.5. The disappearance of the rescaled outlier at *γ* = *f* indicates that the network statistics are consistent with the classical Hebbian process described in Sec. 2.1, demonstrating once again that the system fully recovers flexibility only when inhibition is locally balanced with the fraction of active cells.

Finally, we wondered whether local inhibition would work in a more general scenario when also additional inputs are present, as in Secs. 2.2, 2.3. Until now in this section, we only treated the worst case scenario of freezing, i.e., when additional inputs are not present. Indeed, in this case, the freezing is drastic. The presence of additional, uncorrelated inputs, approximated as noise, naturally reduces output correlations, leading to less freezing. Whatever the degree of freezing, the mechanism of locally balanced inhibition eliminates it, improving performance, see Fig. 6d-f. Indeed, for low level of noise, i.e., in a system-freezing scenario, local inhibition helps prevent freezing, and moreover, the best value for the inhibitory constant *γ* remains proportional to the fraction (*f* = 0.5) of active cells. When setting *γ* = *f*, the correlation *corr*(*z*^*t*^, *z*^*t*+1^) is zero for any value of *σ* (Fig. 6e).

### 2.6 Locally balanced inhibition enhances memory strength

As a step further, we wondered if the inclusion of locally balanced inhibition recovers the network’s effective memory capacity. To this aim, after a period of initialization, the system was probed with the same input *x*^0^, giving a different readout 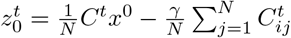 at each time step. As shown in Fig. 7a-b, the correlation 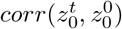 decays in time, for both the classical framework (CF) and the locally balanced inhibition (BI) case, i.e., with *γ* = *f*. Surprisingly, the memory performance for BI improved when compared to CF. This suggests that locally balanced inhibition not only preserves the network’s representational diversity but also extends its operational regime for stable and flexible learning. Note that with this type of models (feedforward network with Hebbian plasticity), forgetting is unavoidable. In fact, in the case of locally balanced inhibition, forgetting is not fully mitigated but indeed postponed.

**Fig. 7:**
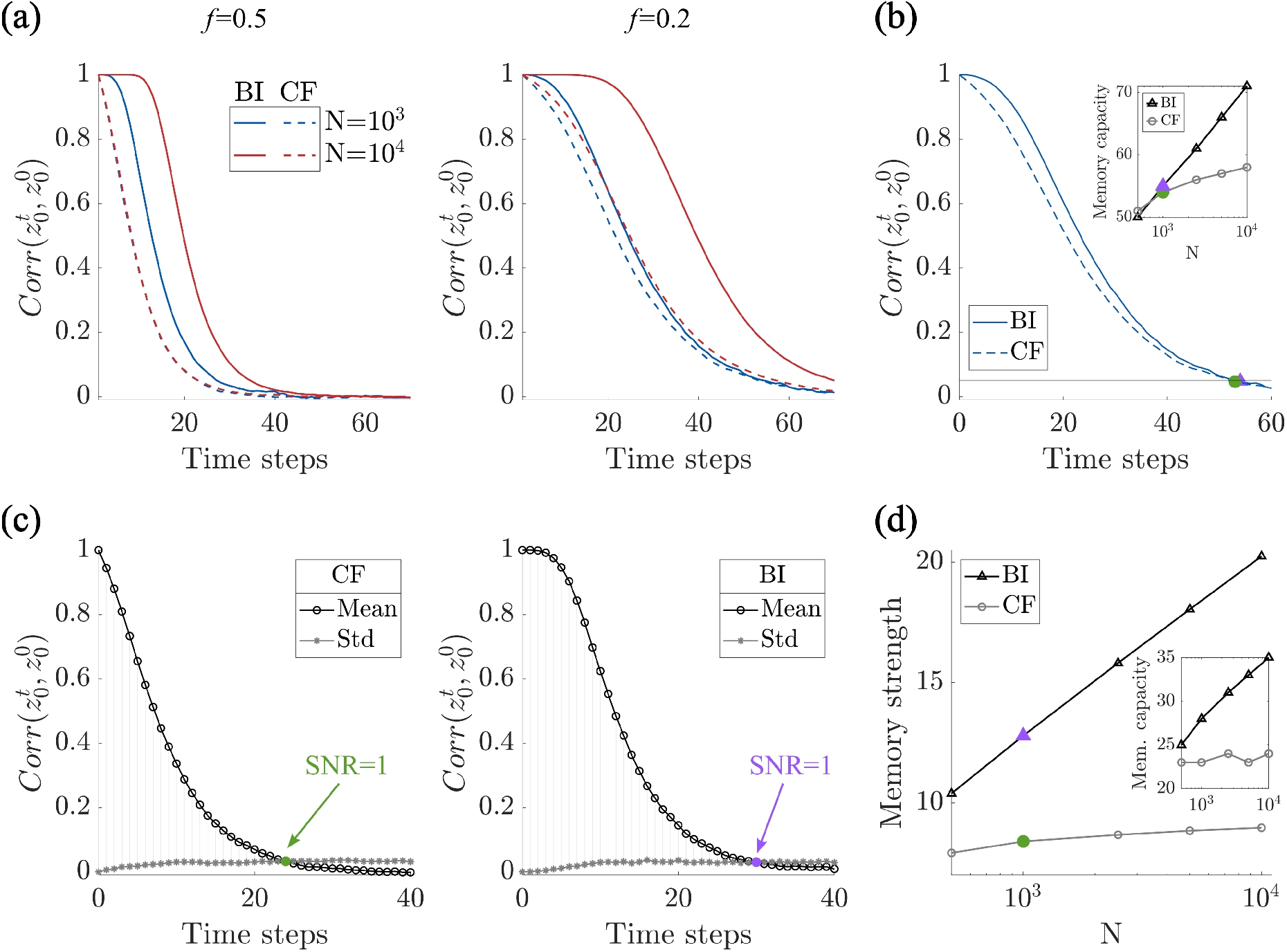
Locally balanced inhibition enhances memory strength. **(a)** The autocorrelation of an elicited output decays over time due to the overwriting from ongoing learning. Such decay is slower for larger *N* and for the locally balanced inhibition case (BI), compared to classical framework (CF). **(b)** Memory capacity, calculated as the number of time steps until the correlation 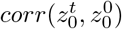 falls below a threshold (0.05) for CF and BI. **(c)** Mean and standard deviation of 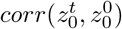 for CF and BI. The color dot identifies the first value of *t* such that the signal-to-noise ratio (SNR) equals 1. **(d)** Memory strength increases logarithmically with the system size, and it does it faster for BI, compared to CF. Inset: Memory capacity metrics for comparison. – Each curve shows the mean over 100 simulations. Parameters: *γ* = *f, f* = 0.5, *p* = 0.2; (b) *f* = 0.2.

To assess the extent to which memory performance improves, we computed the memory strength of the system under these two conditions (CF vs. BI), varying the system size, i.e., the number of neurons *N*. A simple and straightforward metric would be to define memory capacity as the first time step that the correlation 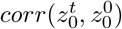 falls below a threshold, see Fig. 7b. The capacity of the BI case is clearly significantly improved compared to the CF. The capacity scales as the logarithm of the system size *N*, a classical finding in both feedforward and recurrent networks with bounded synapses Fusi et al. (2021). This scaling can be improved dramatically by scaling the coding fraction *f* ~1*/N* or by including synapses with a variety of time-scales Fusi et al. (2005); Roxin and Fusi (2013). Apart from this improvement in the memory capacity, the BI case also improves the overall strength of the memory trace, as shown in Fig. 7c. Namely, the correlation does not decay exponentially, as in the CF, but rather remains near one for an extended period of time. To take this effect into account, we defined the memory strength as the area between the mean and standard deviation curves in Fig. 7c, see Fig. 7d.

In summary, locally balanced inhibition serves as an effective mechanism to prevent network rigidity induced by Hebbian plasticity in feedforward architectures. By normalizing synaptic drive at the level of individual neurons, it preserves the diversity of output patterns, and enables the network to store a greater number of input-output associations, while maintaining learning flexibility across time. This highlights the critical role of balanced excitation and inhibition in sustaining functional plasticity within learning systems.

## 3 Discussion and conclusion

In this manuscript, we studied Hebbian hetero-associative networks with stochastic synapses. We first showed how continual learning provokes interference and turnover of synaptic structure, which leads to a decrease in the correlation of driven output patterns over time. We then demonstrated that when the postsynaptic output is driven mainly by a targeted input pathway, i.e., the contribution of additional pathways is negligible, the network enters a pathological freezing regime in which successive readouts become highly correlated. However, even when the contribution of additional inputs is increased, thereby weakening correlations between successive readouts, the targeted pathway still induces output–input correlations, which in turn degrades network performance. To eliminate input-output correlations, we proposed a locally balanced inhibition mechanism, whereby each neuron receives inhibitory input proportional to its own excitatory in-degree. When the inhibitory gain is matched to the network’s activity level, this local balancing recenters the postsynaptic drive. This removes the bias toward repeatedly recruiting the same highly connected neurons and, in turn, decorrelates successive readouts, restoring flexibility in the system. Moreover, in this balanced regime, we observe a clear increase in memory strength.

Classical Hebbian formulations (Hebb (1949); Shatz (1992); Trappenberg (2002)) often assume that the outputs are imposed a priori. By explicitly mixing targeted input drive with additional inputs and varying their relative strength, we bridge imposed-output regimes and biologically motivated input-driven regimes in the same model. Within this continuum, locally balanced inhibition emerges as the minimal circuit-level correction that preserves input-driven readouts while avoiding rigidity, aligning with theoretical (Vreeswijk and Sompolinsky (1996); Murphy and Miller (2009); Renart et al. (2010); Koch et al. (2010); Vogels et al. (2011); Rubin et al. (2015)), and in-vivo (Shu et al. (2003); Haider et al. (2006); Okun and Lampl (2008); Ozeki et al. (2009); Dehghani et al. (2016)) evidence for tight excitation/inhibition balance.

As mentioned, in this manuscript, we introduced a biologically motivated input-driven regime. This necessarily introduces correlations between synaptic inputs and postsynaptic outputs. To the best of our knowledge, prior work on the impact of correlations on Hebbian learning in feedforward architectures has focused on correlations among input patterns and/or among output patterns, not on correlations between inputs and outputs. In (Clopath et al. (2012)), the authors assumed imposed input–output patterns with hand-tuned correlations. In (Földiák (1990); Oja (1982)), to eliminate the correlation between the outputs, it has been proposed to introduce an anti-Hebbian rule between output units. Other works focused on counteracting Hebbian runaway dynamics by distributing changes and maintaining overall balance (Chistiakova et al. (2015); Bannon et al. (2020)). Our work here goes further: in our model, output correlations emerge naturally from uncorrelated inputs, as outputs are driven rather than specified a-priori.

Previous work in recurrent networks has shown that Hebbian plasticity must be accompanied by additional mechanisms, e.g. homeostatic and hetero-synaptic, to allow for the robust formation of neuronal assemblies Zenke et al. (2015); Kalle Kossio et al. (2021). The locally balanced inhibition we consider here could also be interpreted as an additional plasticity mechanism which acts on a time-scale commensurate with or shorter than the time between pattern presentations. Specifically, more active neurons, namely those with greater excitatory drive, would recruit more inhibitory input by strengthening the I to E synapses Vogels et al. (2011). This effectively constitutes a homeostatic mechanism which serves to balance E and I currents. Other homeostatic mechanisms which would succeed in renormalizing input distributions across neurons would also alleviate the freezing we find here, e.g. fixing the total excitatory input to a cell.

Our results connect directly to representational drift, i.e., the gradual reconfiguration of neural codes despite stable behavior (Ziv et al. (2013); Rubin et al. (2015); Khatib et al. (2023); Geva et al. (2023); Devalle et al. (2025); Schoonover et al. (2021)). In the classical Hebbian regime (with weak output–input coupling), continual learning produces turnover of synaptic structure and a slow decay of recall, echoing drift-like dynamics. By contrast, strong coupling collapses drift into freezing: the code no longer reconfigures but instead converges to a stereotyped pattern, impairing memory. Locally balanced inhibition prevents freezing while preserving ongoing variability driven by current inputs, supporting flexible, but not runaway, reorganization of representations. This suggests a circuit-level account of how cortex can exhibit drift without loss of function: balanced E/I suppresses correlation-driven positive feedback, maintaining diversity of readouts and extending the regime where behavior remains stable as codes evolve.

Although our model is a strictly feedforward network with binary states, we expect that using real-valued activities would yield qualitatively similar results. On the other hand, future work should examine how alternative plasticity rules shape the dynamics. In this regard, the recent work of Confavreux et al. (2025), which introduces an automated search algorithm to generate thousands of diverse plasticity-rule quadruplets, offers a useful starting point.

All together, by introducing a mechanism of locally balanced inhibition, we demonstrate a simple yet powerful solution that restores flexibility and preserves memory capacity. These findings emphasize the importance of inhibitory regulation in maintaining the balance between stability and adaptability in neural learning systems.

## 4 Acknowledgement

This project has received funding from Proyectos De Generación De Conocimiento 2021 (PID2021-124702OB-I00) and the Proyecto de Colaboración Internacional PCI2023-145967-2 from the Spanish Ministry of Science and Innovation. This work is supported by the Spanish State Research Agency, through the Severo Ochoa and Maria de Maeztu Program for Centers and Units of Excellence in R&D (CEX2020-001084-M). We thank CERCA Programme/Generalitat de Catalunya for institutional support.

